# Neural oligarchy: how synaptic plasticity breeds neurons with extreme influence

**DOI:** 10.1101/361394

**Authors:** Florence I. Kleberg, Jochen Triesch

**Author notes:** Current Address: Frankfurt Institute for Advanced Studies, Frankfurt am Main, Hessen, Germany.

## Abstract

Synapses between cortical neurons are subject to constant modifications through synaptic plasticity mechanisms, which are believed to underlie learning and memory formation. The strengths of excitatory and inhibitory synapses in the cortex follow a right-skewed long-tailed distribution. Similarly, the firing rates of excitatory and inhibitory neurons also follow a right-skewed long-tailed distribution. How these distributions come about and how they maintain their shape over time is currently not well understood. Here we propose a spiking neural network model that explains the origin of these distributions as a consequence of the interaction of spike-timing dependent plasticity (STDP) of excitatory and inhibitory synapses and a multiplicative form of synaptic normalisation. Specifically, we show that the combination of additive STDP and multiplicative normalisation leads to lognormal-like distributions of excitatory and inhibitory synaptic efficacies as observed experimentally. The shape of these distributions remains stable even if spontaneous fluctuations of synaptic efficacies are added. In the same network, lognormal-like distributions of the firing rates of excitatory and inhibitory neurons result from small variability in the spiking thresholds of individual neurons. Interestingly, we find that variation in firing rates is strongly coupled to variation in synaptic efficacies: neurons with the highest firing rates develop very strong connections onto other neurons. Finally, we define an impact measure for individual neurons and demonstrate the existence of a small group of neurons with an exceptionally strong impact on the network, that arise as a result of synaptic plasticity. In summary, synaptic plasticity and small variability in neuronal parameters underlie a neural oligarchy in recurrent neural networks.

**Author summary:** Our brain’s neural networks are composed of billions of neurons that exchange signals via trillions of synapses. Are these neurons created equal, or do they contribute in similar ways to the network dynamics? Or do some neurons wield much more power than others? Recent experiments have shown that some neurons are much more active than the average neuron and that some synaptic connections are much stronger than the average synaptic connection. However, it is still unclear how these properties come about in the brain. Here we present a neural network model that explains these findings as a result of the interaction of synaptic plasticity mechanisms that modify synapses’ efficacies. The model reproduces recent findings on the statistics of neuronal firing rates and synaptic efficacies and predicts a small class of neurons with exceptionally high impact on the network dynamics. Such neurons may play a key role in brain disorders such as epilepsy.

## Introduction

Are cortical networks “democratic” structures in which the voice of every neuron has about the same weight? Or do some neurons wield much more influence than others? And, if so, what mechanisms give rise to this concentration of power? Recent experiments have suggested large inequalities between neurons and synapses. Specifically, a persistent observation is the presence of skewed, long-tailed distributions in firing rates [1–3] and synaptic efficacies [2,4–9]. In most cases, distributions of neuronal variables take on approximately lognormal [3] or power-law [10,11] shapes. Such strongly skewed distributions may have important implications for computation in neuronal networks, for instance effective signal transmission [12], population burst propagation [13] and enlarging the dynamical range of the network [14]. Since the strength of individual synapses in networks of neurons is under constant change due to activity-dependent and homeostatic plasticity [15,16], as well as spontaneous fluctuations [6,17], it is unclear how long-tailed distributions of synaptic weights arise and remain stable. Recent recurrent neural network models of the cortex have demonstrated that long-tailed weight distributions for excitatory-to-excitatory weights can be achieved by a combination of spike-timing dependent plasticity (STDP) and homeostatic mechanisms or structural plasticity of synapses [18–20]. Furthermore, at a phenomenological level it has been argued that such distributions can result from a simple stochastic model called a Kesten process [21]. Recently, inhibitory weights have also been observed to follow a strongly skewed distribution in the hippocampus [22] and in neuronal cultures of the cortex [23]. Although inhibitory synapses are also subject to STDP (iSTDP; [24–26], and homeostatic processes [27,28], it is currently not known how long-tailed distributions of inhibitory weights arise simultaneously with long-tailed distributions of excitatory weights in the cortex, together with long-tailed distributions of firing rates in excitatory and inhibitory neurons [1]. Although a previous study has proposed learning rules that could connect both properties [29], it is unclear how these distributions arise under the influence of biologically plausible plasticity mechanisms, and how they persist in the presence of spontaneous synaptic remodeling [6,17]. Additionally, the long-tailed weight distribution and long-tailed firing rate distribution might be linked at the neuron level, for example in the form of ‘hub’ neurons. Hub neurons are thought to be highly connected in terms of number of synapses and/or synaptic strength, and exert a strong influence on network activity, [30–33], correlations [34–37], and information propagation [33]. Indeed, neurons with high firing rates may form strongly interconnected populations [38].

In this study, we investigate whether the combination of STDP and synaptic normalisation can explain long-tailed distributions of inhibitory and excitatory synaptic weights and firing rates, and how the two are connected. We also address the contribution of spontaneous activity-independent fluctuations in synaptic efficacies [17].

We investigate these issues using a model from the family of self-organising recurrent neural networks (SORN; [20,39]) with leaky integrate-and-fire neurons, and STDP in excitatory and inhibitory synapses. The proposed model is the first to simultaneously explain 1) lognormal-like distributions of excitatory and inhibitory firing rates from small variability in spiking thresholds, 2) the simultaneous emergence of long-tailed lognormal-like distributions of excitatory-to-excitatory and inhibitory-to-excitatory synaptic efficacies, and 3) their persistence in the face of substantial spontaneous weight remodeling. Furthermore, the model predicts a form of “neural oligarchy” — the existence of a small group of extremely powerful neurons that exert a strong influence over the rest of the network.

## Materials and methods

### Neuron and network model

We adapt the self-organising recurrent neural network model (SORN; [39]) to work with leaky integrate-and-fire (LIF) neurons (LIF-SORN; [20]). Our model consists of 400 excitatory and 80 inhibitory conductance-based LIF neurons. The excitatory-to-excitatory (E-E), excitatory-to-inhibitory (E-I), inhibitory-to-excitatory, (I-E), and inhibitory-to-inhibitory (I-I) connections are initialised randomly with a connection probability of 0.02 for E-E connections, 0.1 for E-I and I-E connections, and 0.5 for I-I connections (Fig. 1A, Table 1). Autapses are not allowed. Each connection type has a fixed delay.

**Fig 1.**
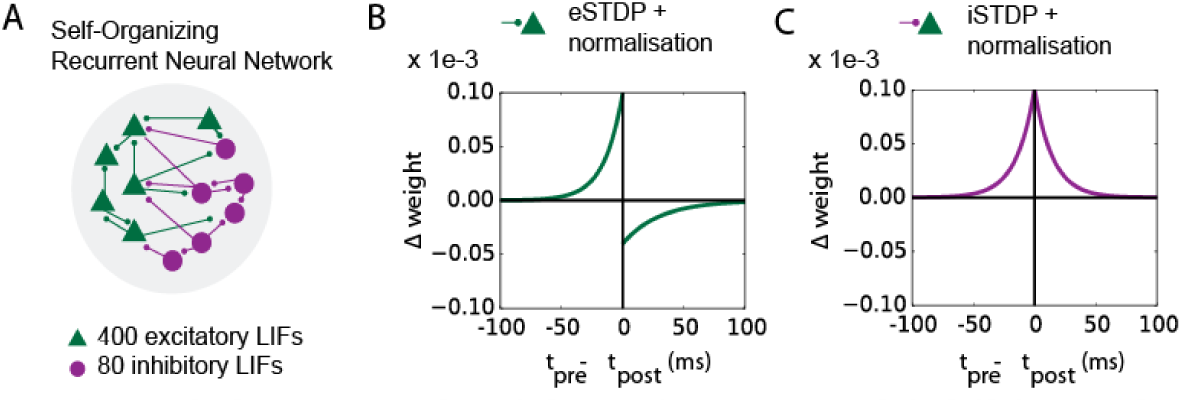
Overview of the network with excitatory and inhibitory STDP. A: Schematic of the network with randomly connected excitatory and inhibitory LIF neurons. B: The STDP learning rule in the E-E weights (eSTDP). The dark green curve shows the amount LTD for a weight at *w*_max,e_. The synapses are also equipped with synaptic normalisation (equation (8) and (9)). C: The STDP learning rule in the I-E synapses (iSTDP).

**Table 1.** Parameters of the network model.

| Network parameters | Description | Value |
| --- | --- | --- |
| $N_e$ | number of excitatory neurons | 400 |
| $N_i$ | number of inhibitory neurons | 80 |
| $p_{e,i}$ | connection probability from excitatory to excitatory neuron | 0.02 |
| $p_{e,i}$ | connection probability from excitatory to inhibitory neuron | 0.1 |
| $p_{i,e}$ | connection probability from inhibitory to excitatory neuron | 0.1 |
| $p_{i,i}$ | connection probability from inhibitory to inhibitory neuron | 0.5 |
| $d_{e,e}$ | axonal delay from excitatory to excitatory neuron | 1.5 ms |
| $d_{e,i}$ | axonal delay from excitatory to inhibitory neuron | 0.5 ms |
| $d_{i,e}$ | axonal delay from inhibitory to excitatory neuron | 1.0 ms |
| $d_{i,i}$ | axonal delay from inhibitory to inhibitory neuron | 1.0 ms |

Table legend: Network parameters.

The LIF membrane equation for a single excitatory or inhibitory point neuron in the network is:

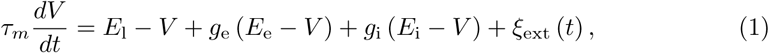

where *V* is the membrane potential in volt, *E*_l_ the leak reversal potential, and *E*_e_ and *E*_i_ are the reversal potentials for excitatory and inhibitory synaptic inputs, respectively. The synaptic conductances are *g*_e_ for excitation and *g*_i_ for inhibition. The values of *τ*_e_ and *τ*_i_ are chosen based on kinetics of AMPA and GABA receptors [40]; see Table 2 for all neuronal parameters. The term *ξ*_ext_ provides an external input of 1 mV to the membrane voltage of the excitatory and inhibitory neurons, at random, Poisson distributed times. Concretely, the external input times for each neuron are sampled from an exponential distribution with *τ*_ext_ = 3 ms, at a resolution of 0.1 ms, resulting in on average 333 input events per second.

**Table 2.** Neuronal parameters.

| Neuronal parameters | Description | Value |
| --- | --- | --- |
| $\tau_e$ | EPSP time constant | 3 ms |
| $\tau_i$ | IPSP time constant | 10 ms |
| $\tau_m$ | membrane time constant | 20 ms |
| $E_e$ | excitatory reversal potential | 0 mV |
| $E_i$ | inhibitory reversal potential | -80 mV |
| $E_l$ | leak reversal potential | -60 mV |
| $\mu_{V_\theta}^e$ | mean firing threshold for excitatory neurons | -50 mV |
| $\mu_{V_\theta}^i$ | mean firing threshold for inhibitory neurons | -51 mV |
| $\sigma_{V_\theta}^e$ | variance of firing threshold for excitatory neurons | 1 mV |
| $\sigma_{V_\theta}^i$ | variance of firing threshold for inhibitory neurons | 1 mV |
| $V_{\text{reset}}$ | reset potential | -60 mV |
| $\xi_{\text{ext}}$ | amplitude of external input | 1 mV |
| $\tau_{\text{ext}}$ | decay constant of exponential sampling of external input times | 3 ms |

Table legend: Neuronal parameters.

For every incoming spike, the conductance of the associated synapse is increased by the value of the weight *w*_e_ or *w*_i_:

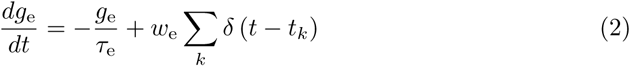

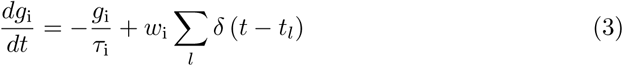

where *t_k_* and *t_l_* are spike times from an excitatory or inhibitory input respectively, and *w*_e_ is the synaptic weight of the excitatory connection and *w*_i_ that of the inhibitory connection. When the membrane potential reaches the threshold *V_θ_*, a spike is generated and the membrane potential is reset to the resting potential *V*_reset_

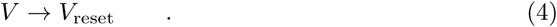

The spiking thresholds *V_θ_* are sampled for each neuron from a Gaussian distribution with mean 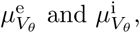 and standard deviation 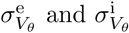 for excitatory and inhibitory neurons, respectively. Doing so creates heterogeneous firing rates in the neuronal population of the model. Membrane potential values are initialised as uniformly random between -50 mV and -55 mV. We set *V*_reset_ = *E*_1_. There is no absolute refractory period for spiking. In equations (1), (2) and (3), conductances are unitless and therefore so are the synaptic weights.

### Synaptic Plasticity

The network incorporates several different types of plasticity. The E-E synapses are endowed with temporally asymmetrical spike-timing dependent plasticity (eSTDP; Fig. 1B), and the I-E connections with symmetrical STDP (iSTDP; Fig. 1C). Both E-E and I-E synapses are subject to homeostatic normalisation of the weights, described in detail below.

Contrary to previous works (e.g. [18,20]) there is no intrinsic homeostatic plasticity and no structural plasticity in this model. This choice was made in order to allow a broad distribution of firing rates to emerge from Gaussian variability in spiking thresholds, and to demonstrate that the STDP rules and the synaptic normalisation are sufficient to account for stable activity levels.

The eSTDP rule is as follows: For the temporal difference between a presynaptic and a postsynaptic spike Δ*t* = *t*_pre_ – *t*_post_, the change in synaptic efficacy is given by:

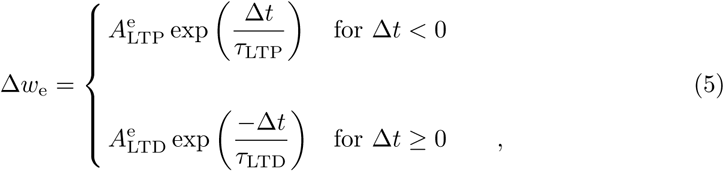

with *A*_LTP_ > 0 and *A*_LTD_ < 0. In this model, eSTDP and iSTDP favour LTP over LTD [41]. The depression area within the eSTDP window is set to 80 percent of the potentiation area, creating here a small advantage for LTP. In synapses from inhibitory to excitatory neurons, additive iSTDP is applied:

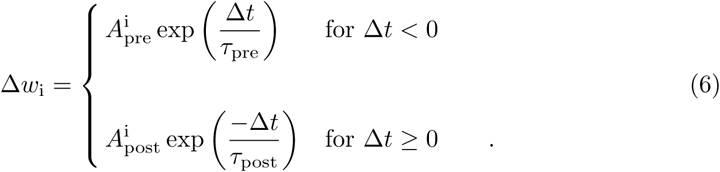

We consider by default *τ*_pre_= *τ*_post_ = *τ*_LTP_, so that the time constants for iSTDP match the LTP time constant in eSTDP. The values of 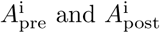 are identical in the symmetric iSTDP learning rule (Fig. 1C). The symmetric iSTDP window, first employed in a recurrent neural network that self-organises into a balanced state [42] has been shown recently to exist in auditory cortex [26]. Besides the symmetric iSTDP (Fig. 1C), we also test a pre-LTP and a post-LTP iSTDP (Suppl. Fig. 4A).

To include a form of depression in the iSTDP, a fixed value LTD_*α*_ is subtracted for every pre-synaptic spike [42]

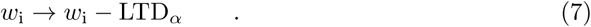

The LTD part of iSTDP can be included by setting LTD_*α*_ to nonzero values. We consider iSTDP with LTD_*α*_ = 0 and with LTD_*α*_ = 0.02. Neither E-E nor I-E weights are allowed to become negative.

Besides eSTDP and iSTDP, the synapses are equipped with a homeostatic mechanism: synaptic normalisation (SN) of the E-E and I-E weights, scaled proportionally to the total incoming weight onto each excitatory neuron. This fast, homeostatic mechanism [43] is based on the premise of having limited numbers of synaptic building blocks such as neurotransmitter receptors or scaffolding proteins in the postsynaptic neuron at any time, which leads to a competition for these building blocks between synapses of the same type [44]. Although there exists evidence for SN in glutamatergic and GABAergic synapses [27,28,43] as well as slow scaling [9,45] within dendritic branches [46], we make the assumption that the SN in these synapses is fast, and that the homeostatic change depends on the weight in a multiplicative fashion [44]. The SN works as follows: all excitatory (inhibitory) weights onto a neuron are regularly updated as

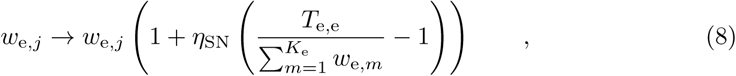

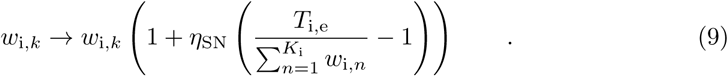

where *T*_e,e_ is the target sum of E-E weights, and *T*_i,e_ the target sum of I-E weights onto one excitatory neuron (see Table 3 for plasticity parameters). The values *K*_e_ and *K*_i_ are the total number of excitatory and inhibitory synapses onto that particular neuron in the network. Since connectivity is initialised at random between neuron pairs, *K*_e_ and *K*_i_ follow a binomial distribution over the population. The E-E and I-E weights are all initialised at 0.0015. The E-I and I-I weights, which are not subject to eSTDP or iSTDP in our model, are initialised by full rescaling so that the weights onto each neuron sum exactly to *T*_e,i_ and *T*_i,i_, respectively (see Table 3 for values). The SN for E-E and I-E weights (equations (8) and (9)) is implemented every second, to save computation time.

**Table 3.** Plasticity parameters.

| Plasticity parameters |  |  |
| --- | --- | --- |
| STDP | Description | Value |
| $A_{LTP}^e$ | amplitude of LTP in eSTDP | $0.1 \times 10^{-3}$ |
| $A_{LTD}^e$ | amplitude of LTD in eSTDP | $-0.04 \times 10^{-3}$ |
| $A_{pre}^i$ | amplitude of iSTDP for $\Delta t < 0$ | $0.1 \times 10^{-3}$ |
| $A_{post}^i$ | amplitude of iSTDP for $\Delta t > 0$ | $0.1 \times 10^{-3}$ |
| $\tau_{LTP}$ | decay constant of LTP in eSTDP | 15.0 ms |
| $\tau_{LTD}$ | decay constant of LTD in eSTDP | 30.0 ms |
| $\tau_{pre}$ | decay constant of iSTDP for $\Delta t < 0$ | 15.0 ms |
| $\tau_{post}$ | decay constant of iSTDP for $\Delta t > 0$ | 15.0 ms |
| $LTD_{\alpha}$ | pre-spike triggered LTD for iSTDP | $0.0 \sim 0.02 \times 10^{-3}$ |
| Synaptic Normalisation | Description | Value |
| $T_{e,e}$ | max input sum of E-E synapses | $50 \times 10^{-3}$ |
| $T_{e,i}$ | max input sum of E-I synapses | $60 \times 10^{-3}$ |
| $T_{i,e}$ | max input sum of I-E synapses | $15 \times 10^{-3}$ |
| $T_{i,i}$ | max input sum of I-I synapses | $60 \times 10^{-3}$ |
| Spontaneous Weight Changes | Description | Value |
| $\mu_e$ | mean of spontaneous E-E weight changes | $0.025 \times 10^{-3}$ |
| $\mu_i$ | mean of spontaneous I-E weight changes | $0.025 \times 10^{-3}$ |
| $\sigma_e$ | standard deviation of spontaneous E-E weight changes | $0.1 \times 10^{-3}$ |
| $\sigma_i$ | standard deviation of spontaneous I-E weight changes | $0.1 \times 10^{-3}$ |

Table listing all plasticity parameters. Although E-I and I-I synapses are not subject to plasticity during the simulation, $T_{e,i}$ and $T_{i,i}$ are used at network initialisation to set the weights for these connections.

The simulations of the network run on custom-built code in Python and the neural simulator Brian [47].

### Quantification of spike correlations and firing rates

To compute the spike-time correlations between neurons, we bin the spikes in 100 ms bins. Since spike-time correlations depend on the bin size for the spikes [48], we choose the bin size large enough to allow for large enough correlations, but so as not to exceed the mean inter-spike interval for excitatory and inhibitory neurons [48]. To obtain the firing rates for each neuron, the total number of spikes in each neuron is simply divided by the length of the entire simulation. The histograms of firing rates that are shown on a semilog scale contain exponentially distributed bins, resulting in equally spaced bins in the figure. These histograms are normalised by dividing each bin value by the bin width, as was done in previous work highlighting lognormal distributions [5]. A lognormal curve is fitted to the distributions using the SciPy package curve_fit. The histograms on linear axes are not normalised unless stated explicitly. We investigate if the firing rate distribution in the population is a direct result of the interaction between the distribution of membrane potentials of the neurons, and their spiking thresholds. For this we acquire the distribution of membrane potentials visited over 50 seconds by a neuron in which spiking is disabled, and average over *N*_e_ of such distributions. The expected number of spikes for each neuron is then computed as the area of the mean membrane potential distribution for a given spiking threshold sampled from the Gaussian distribution. This process is repeated for 10000 putative neurons.

### Analysis of synaptic efficacy distributions

We record the weights in the network at every second, just before and just after SN is applied. We verify if the weight dynamics reaches an equilibrium by checking if the mean and standard deviation of the weight distribution reach a stable point over time. We then create histograms of the weight distribution at the final time point of the simulation, after the distributions of E-E and I-E weights have stabilised (40000 seconds). The visualisation of the weight distribution follows the same rules as the firing rate distribution.

### Definition of hub neurons

Hub neurons are neurons with stronger and more numerous outgoing connections. Since the number of outgoing connections is initialised randomly and fixed during the simulation, only hub neurons with stronger connections are considered here. To detect the presence of hub neurons, we split the excitatory neurons into a group with the ten percent highest firing rates (high) and the rest of the neurons (control). The outgoing weights from the high group and the control group are then compared: if the high group has significantly larger outgoing weights on average, this suggests that the high-firing neurons are also better connected. The inhibitory neurons are compared in the same way. Only the plastic synapses (E-E and I-E weights) are taken into account. To determine the impact of a neuron *m* on the network, we employ

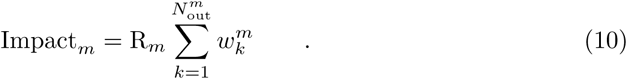

where R_*m*_ is the average firing rate of the neuron recorded over 200 seconds after weight stabilisation, 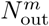 the number of its efferent synapses, and 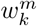 the weight of each of these synapses at the end of the 200 second interval. The impact is verified for excitatory neurons with their E-E synapses, and inibitory neurons with their I-E synapses.

### Stochastic models of weight dynamics

We hypothesise that the weight distributions found in the network simulation can be explained by simple stochastic processes. We consider three such processes: a uniform stochastic model (USM), a nonuniform stochastic model (NSM), and a Kesten model [21,49]. In these models, a synaptic weight is reduced to a stochastic variable *w* that evolves under random additive and multiplicative changes over a number of steps. First, in the USM we let *w* change according to SN and STDP, where the STDP changes are random and any spike-time combination is assumed to be equally probable. Therefore, the USM disregards the likelihood of specific correlations between pre- and post spike times at a synapse. We start with a random variable *w_t_* that represents a synaptic weight at a time step *t*. We keep I-E weights and E-E weights in the stochastic models between 0 ≤ *w_t_* ≤ *T*_i,e_, and 0 ≤ *w_t_* ≤ *T*_e,e_, respectively. The weight *w_t_* is modified *N*_step_ = 5000 times in an additive way by randomly sampling an STDPupdate 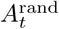 from the STDP windows shown in Fig. 1B-C. However, some neurons in the network have intrinsically higher firing rates compared to other neurons, which may give some weights an advantage over others when competing for limited postsynaptic resources. We therefore assume that some weights benefit from more frequent STDP updates. We sample the number of updates λ_STDP_ for each *w^m^* from a lognormal distribution that is fitted to the observed firing rates in the network. The number of updates 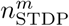 for each 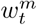 is sampled from a Poisson distribution with mean 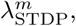 since the average firing rate in each neuron and thus for each weight in the simulation does not change over time but the number of updates in each step is variable. For each time step, a weight *w^m^* therefore receives the update

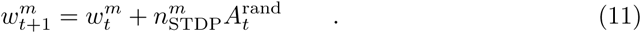

Furthermore, in the network simulation, not all neurons receive the same number of inhibitory or excitatory synapses. How much a weight is changed by SN depends on the number of competing neighbours it has. We therefore mimic the variability in in-degree of I-E (E-E) synapses by sampling a number *n*_total_ from a binomial distribution, where *n*_total_ represents the number of inhibitory or excitatory synapses onto a postsynaptic excitatory neuron given the random connectivity, numbers of neurons, and connectivity fractions in the network for inhibitory-to-excitatory and excitatory-to-excitatory neurons. We take *n*_total_ instances of *w*^m^ and normalise them together with equation (8) and (9). Normalisation is performed at every step, although the same results are obtained when SN is slower. The process is then repeated *N*_e_ times, to represent all postsynaptic excitatory neurons in the network. The total distribution of *w* should then closely resemble the distribution of weights in the network if its shape is purely the result of random STDP updates combined with multiplicative SN.

Second, we consider a nonuniform stochastic model (NSM), which is identical to the USM in all aspects but one: the STDP updates are not sampled randomly from the STDP windows, but the sampling probabilities are weighted by the spike-time cross-correlations between neuron pairs. To obtain these correlations, spikes are recorded from the network for 200 seconds after the weights have reached equilibrium, and cross-correlations are computed as Peason’s correlation coefficients over spike-time lags between -100 and 100 ms surrounding the spike, covering the relevant part of the eSTDP and iSTDP windows, using spike bins of 5 ms. These correlations are averaged over all connected I-E and E-E neuron pairs, and averaged over 10 trials. Cross-correlation curves are then shifted upwards additively by the minimal value to remove negative values, normalised to obtain an area of 1.0, and overlaid with the STDP windows to weigh sampling probabilities for the NSM.

Third, we consider that the weights might behave according to a Kesten process [49]. A variable *w* that follows a Kesten process is defined by the following iteration:

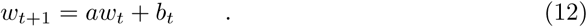

where *a* and *b* are independent, stochastic variables. Although the Kesten process can be seen as analogous to the combination of additive STDP and multiplicative SN [50], in our model STDP and SN are not strictly independent nor stochastic, since both terms depend on the collective behaviour of groups of weights and their current state. We can however quantify the difference of our results from a Kesten process by assuming weighted random STDP sampling as in the NSM, setting 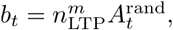 and gleaning the distribution for *a* from the SN factors that were obtained from the network simulation. In the network model, the USM and the NSM, the SN is stabilising the weights in a non-random manner that depends on the collective weight state (Equation (8) and (9)) and therefore also the state of the weight itself. Because of the independent sampling of *a* in the Kesten process, the SN factor is not directly related to the exact value of *w* at any time, as opposed to the USM and NSM. All three stochastic models (USM, NSM, Kesten) are performed for parameters representing both E-E and I-E weights and their distributions are compared to their respective distributions from the network simulations.

### Spontaneous fluctuations of weights

To investigate whether the lognormal-like distributions persist in the presence of spontaneous fluctuations of the synapses, we add random Gaussian noise to the synaptic weights (“weight noise”) at each second, with mean *μ*_we_ = *μ*_wi_ = 0.025 × 10^−3^ and standard deviation *σ*_we_ and *σ*_wi_ = 0.1 × 10^−3^ for for E-E weigths and I-E weights, respectively. Since we consider fluctuations in the order of activity-dependent changes through STDP [17], we set *σ*_we_ and *σ*_wi_ equal to 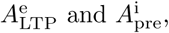 the amplitude of LTP in the eSTDP and iSTDP rules. The weight noise is applied independently in each weight in the network. During the spontaneous fluctuations, we maintain eSTDP (iSTDP) with SN in the E-E (I-E) weights. We also study the effect of TTX on the fluctuations of synaptic efficacies. In the TTX condition, eSTDP and iSTDP are rendered inactive but SN and the spontaneous fluctuations remain active.

## Results

### The network maintains asynchronous irregular activity and low correlations

The LIF-SORN with iSTDP displays a number of properties commonly seen in cortical networks (Fig. 2). Specifically, the spike-correlations are low [51], and spike patterns are irregular for excitatory and inhibitory neurons [52,53]. A sample of population spiking is shown in Fig. 2A, with excitatory (inhibitory) spikes shown in green (purple). Evidently, some neurons spike often and others are relatively silent. In Fig. 2B, a random excitatory neuron receiving excitatory and inhibitory inputs is shown with its membrane potential, excitatory and inhibitory conductances, and currents. To determine whether the neurons spike asynchronously, we calculate the distribution of coefficients of variation (CVs) of inter-spike intervals (ISIs) of each neuron in the network. A CV of 1.0 is a property of a neuron firing according to a Poisson process, while a lower value indicates more periodic firing patterns. Figure 2C shows that both excitatory neurons (mean of distribution: 0.77) and inhibitory neurons (mean of distribution: 0.76) fire moderately aperiodically. Moreover, spike-time correlations between excitatory neurons, between inhibitory neurons, and between excitatory and inhibitory neurons are low, indicating a network with low synchrony (Fig. 2D). Though the network activity is not perfectly decorrelated, based on these observations we conclude that the network is close to the asynchronous state [54,55].

**Fig 2.**
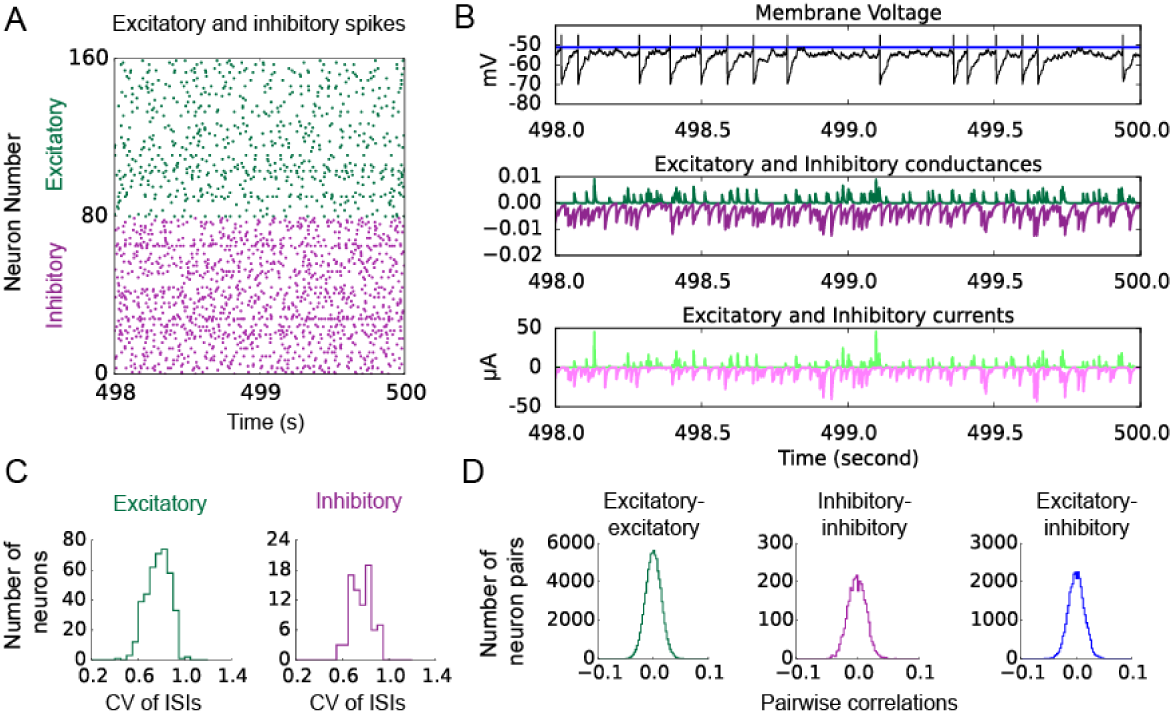
Activity in the network with eSTDP, symmetric iSTDP, and SN. A: Spike raster plot of the inhibitory population, and a portion of the excitatory population, during the 498th to 500th second of a simulation. B: A momentary view of the membrane potential and synaptic currents onto a representative excitatory neuron in the network during the same interval as in A. Top, membrane potential. The firing threshold is indicated in blue. Spikes are shown with vertical lines. Middle, conductance dynamics (see equations (1), (2), (3)). Bottom, excitatory (green) and inhibitory (red) currents. C: Coefficient of variation of the inter-spike intervals of excitatory (left) and inhibitory (right) neurons, during a 500 second simulation. D: Distributions of spike cross-correlations between neurons (left, excitatory-excitatory; middle, inhibitory-inhibitory; right, excitatory-inhibitory) taken from a 500 second simulation. Correlations were computed using 100 ms bins for the spikes.

### Development of lognormal-like distributions of firing rates

Since the incoming excitatory and inhibitory conductance onto neurons in the network is normalised over time (equation (8) and (9)), setting all neuronal parameters equal leads to a network where all excitatory or inhibitory neurons fire on average at the same rate. However when we sample the spiking thresholds of individual neurons from a normal distribution with 1 mV variance, the average firing rates in the network become heterogeneous across neurons. Importantly, the distributions of firing rates of excitatory and inhibitory neurons follow a lognormal-like shape (Fig. 3A-B) as seen in recordings from rat auditory cortex [1] and hippocampus [2]. Because it is not directly clear how lognormal-like distributed firing rates relate to normally distributed spiking thresholds, we propose that the observed firing rates are directly the result of the interaction between the spiking threshold sampling distribution (Fig. 3C, red curve) and the values that the membrane potential in an average neuron typically visits over time, when its spiking mechanism is disabled (Fig. 3C, blue curve). The expected number of spikes can then be computed by iteratively sampling from the spiking threshold curve and determining how many times the average neuron’s membrane voltage crosses the threshold. The resulting distributions of expected number of spikes per neuron follow a lognormal-like shape (Fig. 3D), suggesting that lognormal-like distributed firing rates can already arise from the interaction of near-Gaussian membrane dynamics and small Gaussian variability in the spiking thresholds.

**Fig 3.**
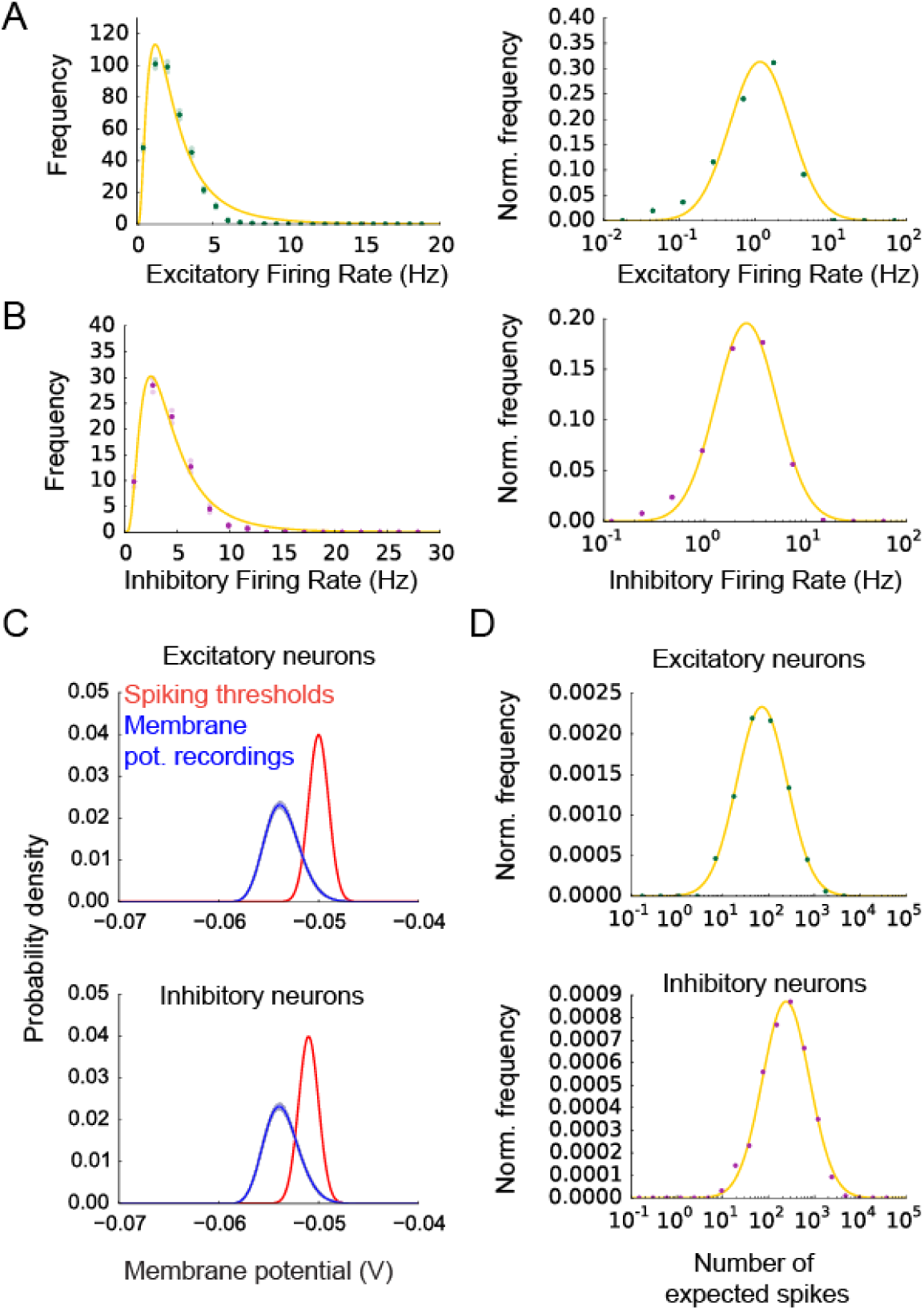
Lognormal distributions of firing rates in excitatory and inhibitory neurons. A: Mean distributions of firing rates of the excitatory neurons during simulations of 5000 seconds. Firing rates are obtained from the total number of spikes during the simulation intervals. The light colour shows the standard error. The yellow curves show lognormal fits. The distributions are shown both on a linear and a semilog scale and are averaged over 10 trials. In the bottom figure, the histogram binsizes are corrected to be linear on the logscale, and bin values are corrected by dividing them through the bin width. B: Same as in A but for inhibitory neurons. C: Blue, mean distributions of membrane voltage recordings from 400 neurons over 50 seconds, where spiking is disabled. The light blue indicates the standard deviation over neurons. Red, Normal distribution used for sampling spiking thresholds at network initialisation. D: Distribution of expected number of spikes estimated for 10000 neurons, based on sampling from the blue and red curves in C. Top, excitatory neurons. Bottom, inhibitory neurons.

### Lognormal-like excitatory and inhibitory weight distributions

Next, we wish to find how synaptic weight distributions are shaped by plasticity mechanisms. During self-organisation, the E-E (I-E) weights evolve under the influence of eSTDP (iSTDP) and SN. An example of the dynamics of weights during the simulation is shown in Fig. 4A-B. Due to the SN in E-E and I-E synapses, the mean weight in the network remains approximately constant, after some initial transients which are due to ‘soft’ normalisation (equations (8) and (9); Fig. 4A-B, thick lines). After 40000 (10000) seconds of self-organisation, the standard deviation of the E-E weight (I-E weight) distribution stabilises (Suppl. Fig. 1). The weights have formed stable, right-skewed long-tailed distributions (Fig. 4C-D; a lognormal fit is shown in the orange curve, see also videos ?? and ??), although the individual E-E and I-E weights continue to change. The weight distributions of the E-E and I-E weights resemble lognormal-like distributions seen in experiments [2,4,5,7–9,23]. In Fig. 5, the mean of the distributions over 10 independent instances of the network is shown. The lognormal-like shapes are still present after 40000 seconds of self-organisation, just before (Fig. 5) and just after (Suppl. Fig. 2A-D) normalisation is applied. The distributions are not exactly lognormal (compare lognormal fit to data points in Fig. 5), in line with experimental findings [5] .

**Fig 4.**
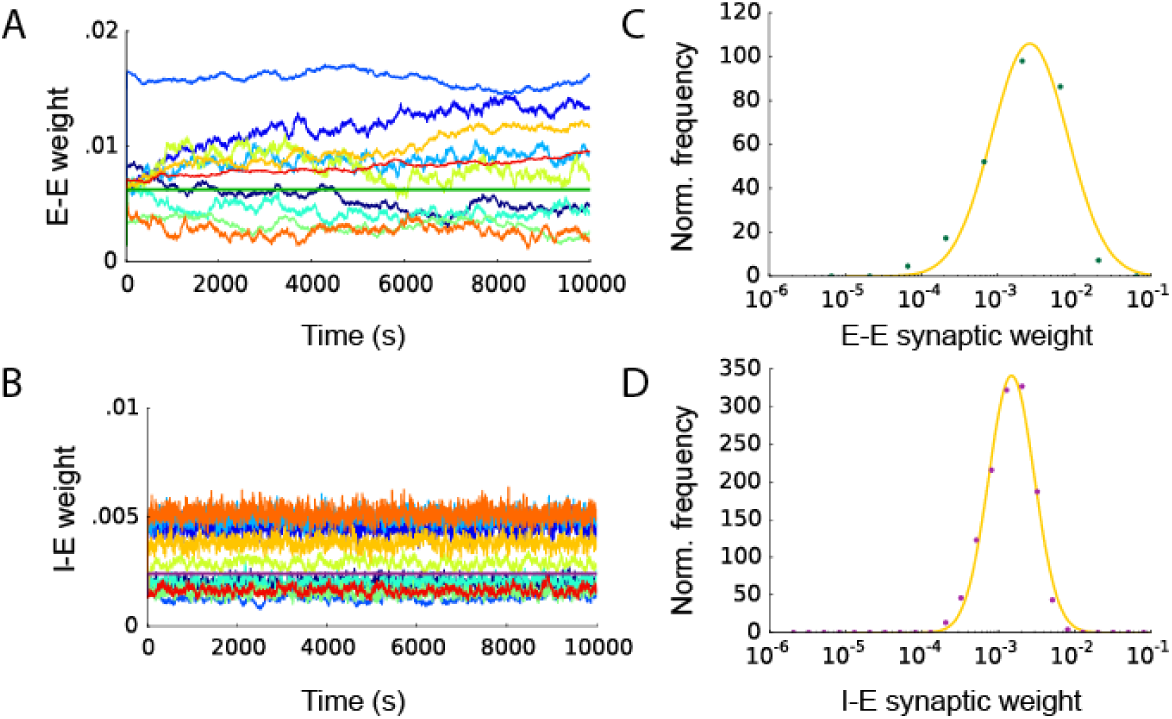
Weight dynamics of a single simulation in the network with eSTDP, iSTDP and SN. A: Evolution of 10 random E-E weights after 10000 seconds of self-organisation. The thick line indicates the mean of all E-E weights in the network. B: Distribution of the E-E weights in a single simulation after 10000 seconds of self-organisation. The orange curve is a lognormal fit. The histogram bins are chosen to be linear on the logscale, and bin values are divided by their corresponding bin width. C: Evolution of 10 random I-E weights. Here, LTD_*α*_ = 0.0. D: Distribution of the I-E weights after 10000 seconds of self-organisation.

**Fig 5.**
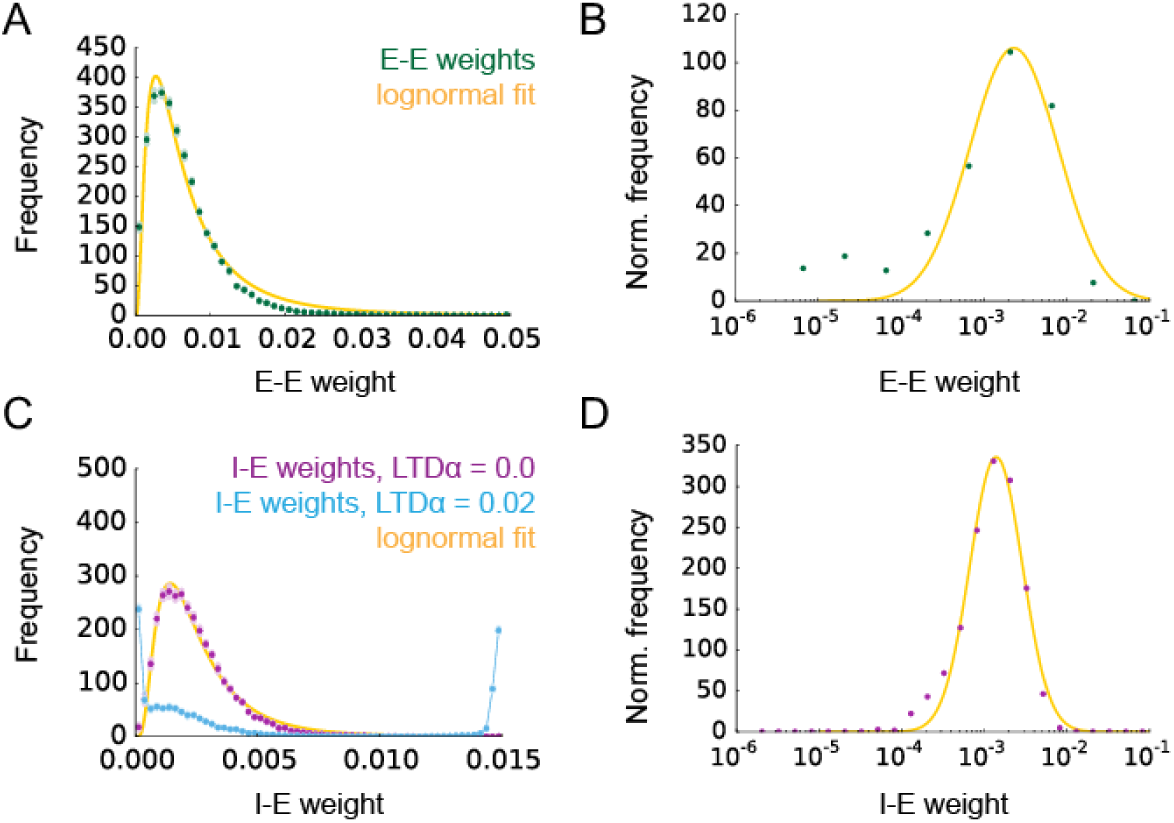
Average distributions of 10 trials of E-E and I-E weights in the network with iSTDP with SN in I-E synapses. A: Mean distribution of E-E weights. The dark point show the average, the light point show the standard error of the mean over 10 independent trials. The orange curve is a lognormal fit to the mean distribution. B: Mean distribution of E-E weights plotted with uniform bin spacing on a logscale, in which bin values are corrected for their corresponding bin width. C: Mean distributions of I-E weights, for LTD_*α*_ = 0 (purple points), and LTD_*α*_ = 0.02 (blue points). A line is drawn between the blue points for visibility. D: As in B but for I-E weights, with LTD_*α*_ = 0. In all subfigures, the weights are recorded just before SN.

For E-E weights, a lognormal-like distribution is found when combining eSTDP with SN (Fig. 5A-B), as was shown before in [20] and [18], in which eSTDP and SN were complemented with structural plasticity. We here use a different approach, leaving out structural plasticity, but applying a small bias in the eSTDP rule, in favour of potentiation. The result is a lognormal-like distribution of E-E weights (Fig. 5A-B). The I-E weights also follow this distribution when LTD_*α*_ = 0 (Fig. 5C-D), meaning the iSTDP is purely potentiating. Thus, the combination of an additive potentiating STDP rule, together with SN results in lognormal-like distributions of E-E and I-E weights. What happens if the STDP rule contains more LTD than LTP, or when SN is left out? When LTD_*α*_ is increased to 0.02, the I-E weight distribution changes and loses its lognormal-like shape (Fig. 5C, blue curve), indicating that strong LTD in the iSTDP rule can move the shape of the weight distribution away from the unimodal shape, pushing most weights to the zero boundary and a small number of weights to the maximum weight. A graphical representation of how LTP and LTD shape the distribution of E-E and I-E weights is shown in Fig. 6. The multiplicative SN pushes competing weights toward each other when LTP is dominant (Fig. 6A), which for a population of weights will lead to a unimodal distribution. Conversely, SN pushes the weights away from each other when LTD is dominant (Fig. 6B), which for a population of weights will lead to a bimodal distribution. Therefore, the lognormal-like shape of the I-E weight distribution observed in Fig. 4 and 5 is a specific result of the interaction of potentiating iSTDP and multiplicative SN. On the other hand, when SN is removed, the weights are allowed to move freely between the zero boundary and the maximal weight boundary through STDP. However, in this case, the distribution becomes distorted and weights heap up at the maximal weight boundary (Suppl. Fig. 2E, purple curve). Likewise, when LTD is added, the distribution shifts leftward again (shown for I-E weights, Suppl. Fig. 2E, blue curve). In other words, when SN is left out, or LTD is too strong, the weight distribution loses its lognormal-like shape, and becomes bimodal.

**Fig 6.**
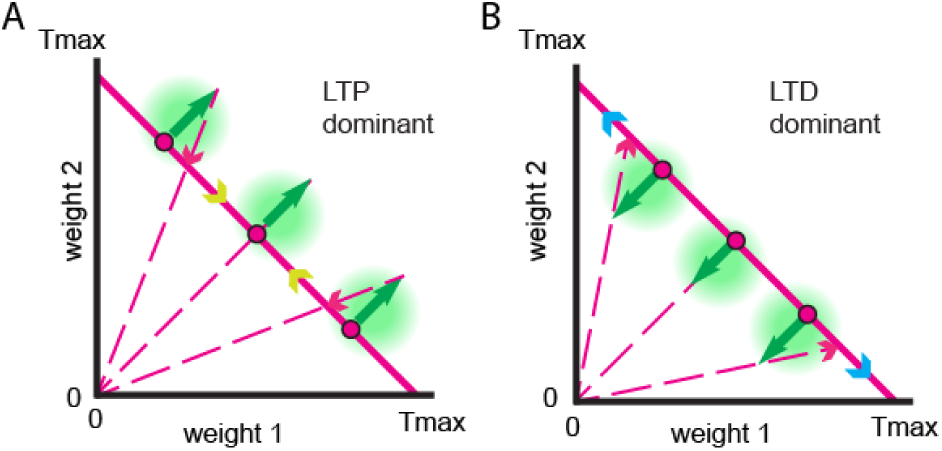
Weight dynamics shaped by LTP, LTD and SN. A: Graphic description of changes in two competing postsynaptic weights under LTP-dominated plasticity (green areas) with multiplicative SN (dotted lines). The sum of the weights is maintained at *T*_max_. When LTP dominates, the two weights are pushed towards the centre, obtaining intermediate values close together. B: An excess of LTD (green areas) pushes two competing weights away from each other through SN (dotted lines), with one weight approaching zero and the other weight gaining nearly all of the *T*_max_.

Lognormal-like distributions have been observed in the weights of cortical [5,8] and hippocampal [2] populations, although the observed distributions usually show slight deviations from a lognormal shape (see also [18,23,56]). We conclude that I-E and E-E weights in our network may form through the combination of potentiating STDP and SN a right-skewed long-tailed distribution that resembles a lognormal distribution, but as in the experimental observations, is not in fact precisely and entirely lognormal.

### Weight-dependent changes through STDP maintain strong weights

To understand how the changes in STDP can lead to such lognormal-like distributions, we investigated how changes in synaptic efficacies depend on the current synaptic efficacy. For the E-E weights, eSTDP contributes both LTP and to a lesser degree, LTD (Fig. 7A, top), following the eSTDP learning rule (Fig. 1B). For the I-E weights, there is only potentiation through iSTDP as we set here LTD_*α*_ = 0 (Fig. 7B, top). For both E-E and I-E weights, the SN provides the downscaling to compensate for the potentiation through eSTDP and iSTDP (Fig. 7A-B, middle). Interestingly, one can see that although the magnitude of downscaling is larger for some large weights, other large weights are not scaled down as much despite their size (observe the blue points that lie close to the zero change line in Fig. 7A-B, middle figures). If all weights in a population are subject to multiplicative weight-dependent depression with the same factor, weights settle into a symmetric distribution [57, 58], but a sublinear dependence of depression on weights can lead to asymmetry in the weight population, through symmetry breaking [59,60]. A sublinear dependence of LTD on weight for larger weights has already been included in an STDP-based learning rule engineered to produce lognormal-like distributions of weights, by maintaining the large weights inside the tail of the distribution [60]. Indeed one can observe in the total changes for each synapse, namely the sum of changes from STDP and SN, that many large weights do not demonstrate absolute changes as large as their smaller counterparts (Fig. 7A-B, bottom figures, compare left to right half of the figure). The same small population of large weights therefore maintains the tail of the lognormal-like weight distribution over time. Such persistently large weights are in line with findings of higher stability of large dendritic spines compared to small spines [6,9,61,62].

**Fig 7.**
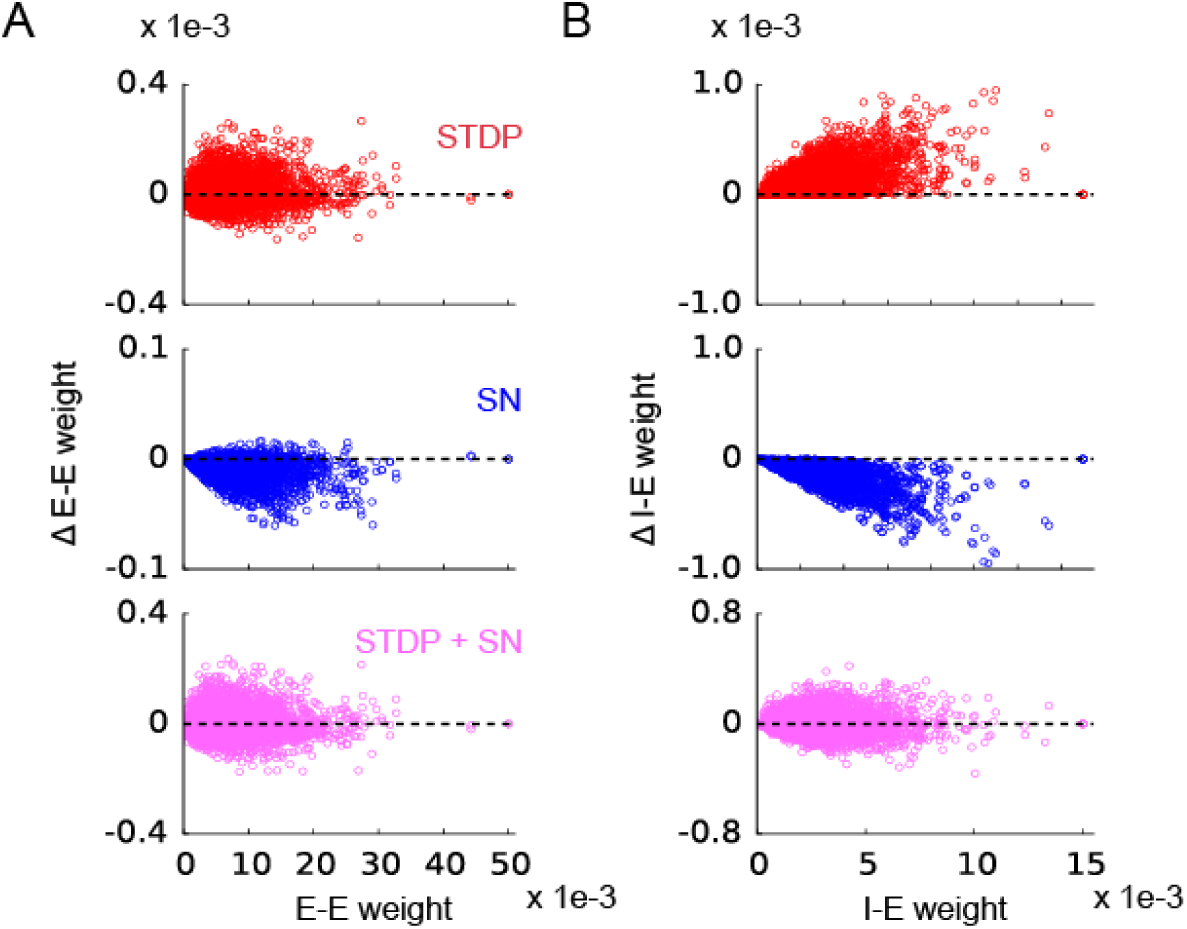
Distribution of synaptic weight changes, as a function of current weight. A: Weight changes for E-E weights. Top, the changes in E-E weight due to eSTDP. Each point represents a separate weight that changed. The dotted line indicates zero change. Changes are recorded during the last two seconds of a 10000 second simulation. Middle, changes in the same E-E weights due to SN. Bottom, combined effects of eSTDP and SN for the same weights. B: Same as in A but for I-E weights and iSTDP with LTD_*α*_ = 0.

### Emergence of excitatory and inhibitory hub neurons

Next, we attempt to find out whether hub neurons are present in the network. We hypothesized that neurons with high firing rates may develop strong outgoing weights due to STDP. Since neurons with high firing rates give rise to more STDP events, their outgoing weights will have an advantage when competing with synapses from less active neurons onto the same target neuron. This principle works well for classical eSTDP rules, but is even more likely to apply when the STDP rule is in itself mostly potentiating (see Fig. 7 and Methods). To investigate the relation between individual neuron’s firing rates and their outgoing weights in our network, we separate the neurons into two groups: the 10 percent highest firing neurons (high-firing), and the remainder of the neurons (control), and observe the mean outgoing weight for each category. Fig. 8 shows that despite considerable variability, the mean outgoing weight of high-firing neurons is larger than that of control neurons. The result is significant for both excitatory neurons with their outgoing E-E weights (Wilcoxon rank-sum test, p= 2.60 × 10^−126^, Fig. 8A), and inhibitory neurons with their outgoing I-E weights (Wilcoxon rank-sum test, p= 1.48 × 10^−121^, Fig. 8B). As all weights start at the same value, the split in the outgoing weight populations happens over time due to plasticity. A neuron with a high firing rate therefore has a higher probability of making its outgoing weights larger, exploiting STDP to achieve a stronger influence on the network. With the above, we replicate findings by Effenberger and colleagues (‘driver neurons’, [19]), and extend them to inhibitory neurons. Moreover, the impact of a neuron on the rest of the network should be a function of both its firing rate and its outgoing weights (see Methods). The distribution of impacts is more strongly skewed and close to exponential, for both excitatory and inhibitory neurons (Fig. 8C-D). In summary, firing rates and STDP-driven competition in synapses combine to create a neural oligarchy with a small number of highly influential excitatory and inhibitory neurons.

**Fig 8.**
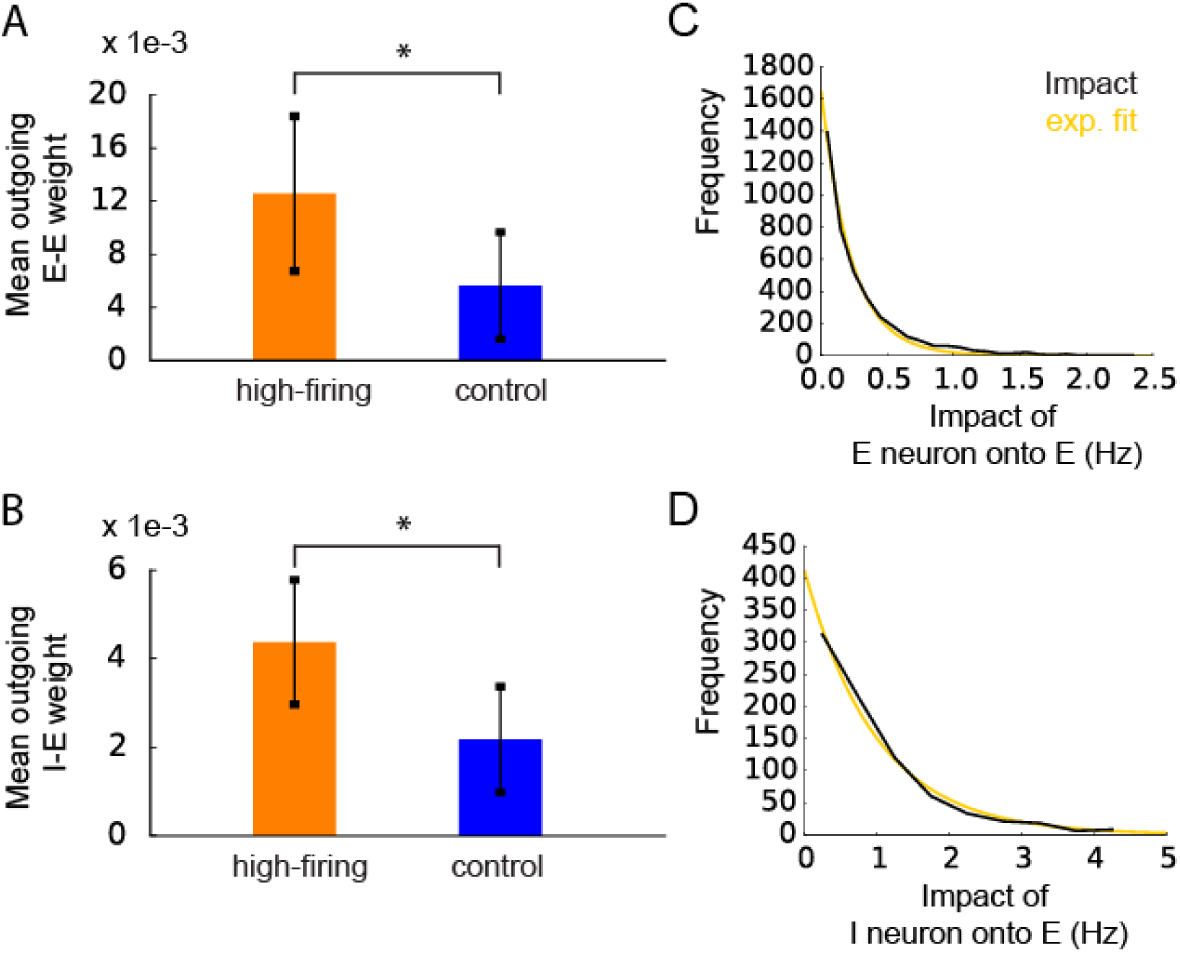
Connection between firing rates and mean outgoing nonzero weight in excitatory and inhibitory neurons. A: Mean outgoing weight for excitatory neurons, sorted into high- and low firing categories (high and control, respectively). The errorbar shows the standard deviation over all weights within the high group and the control group. Data is from a single trial. B: Same as in A but for inhibitory neurons. C: Distribution of impact values for excitatory neurons. The histogram contains neurons from 10 trials of 200 seconds each, recorded after weight stabilisation. The yellow line shows an exponential fit. D: as in C but for inhibitory neurons.

### A simplified stochastic model for excitatory and inhibitory weight dynamics

Since previous studies suggested that a stochastic process can be a good model for the dynamics of synapses [21,50], we investigate whether simplified stochastic models can account for the lognormal-like weight distributions we have shown. In these models, a weight is treated as a stochastic variable *w* that evolves over a number of time steps. We consider stochastic processes which assume the variable *w* is subject to random additive updates sampled from the STDP window (Fig. 1B-C), and collective multiplicative SN (see Methods). We test three such processes, a nonuniform stochastic model (NSM), a uniform stochastic model (USM), and a Kesten model [21,49].

First we consider the NSM, in which *w* is updated by randomly sampling the STDP window in a nonuniform manner. Sampling probabilities in the STDP window are determined by the average correlation coefficient between pre- and postsynaptic spike times, ranging over the relevant time lags for STDP, which is shown in Fig. 9A and B. The distribution of the random variable *w* from the NSM (orange curve), that uses this cross-correlation, closely resembles the mean distribution of E-E weights (Fig. 9C, green curve) and I-E weights (Fig. 9D, purple curve) in the network. This means that the lognormal-like distributions found here are not strongly dependent on variabilities in spike correlations across weights or a specific order of updates by STDP at each weight. Rather, random independent STDP events in the synapses combined with SN govern the qualitative properties of the distributions of E-E and I-E weights to create a lognormal-like shape. The combination of a random additive update with a multiplicative scaling of a stochastic variable as described in the NSM above is reminiscent of a Kesten process, a simple stochastic process that depends on two independent random variables, one additive and one multiplicative [49]. Indeed, a recent study phenomenologically described the dynamics of synaptic weights in networks of cultured neurons with such a process [21]. The Kesten model is here identical to the NSM, except that the multiplicative factor *a* is a random parameter and does not depend on the state of *w*. If we sample from the distribution of SN factors from the network simulation to generate values of *a* and implement a Kesten model for *N*_step_ steps, the Kesten distributions take on a lognormal-like shape (Fig. 9C-D, blue curve), but are not as close to the distributions of E-E and I-E weights from the network simulations (Fig. 9C-D, purple and green curves) or the NSM (Fig. 9C-D, orange curves). Based on the above results, we find that the NSM is a more accurate description of the dynamics of E-E and I-E weights in the network than the Kesten model.

**Fig 9.**
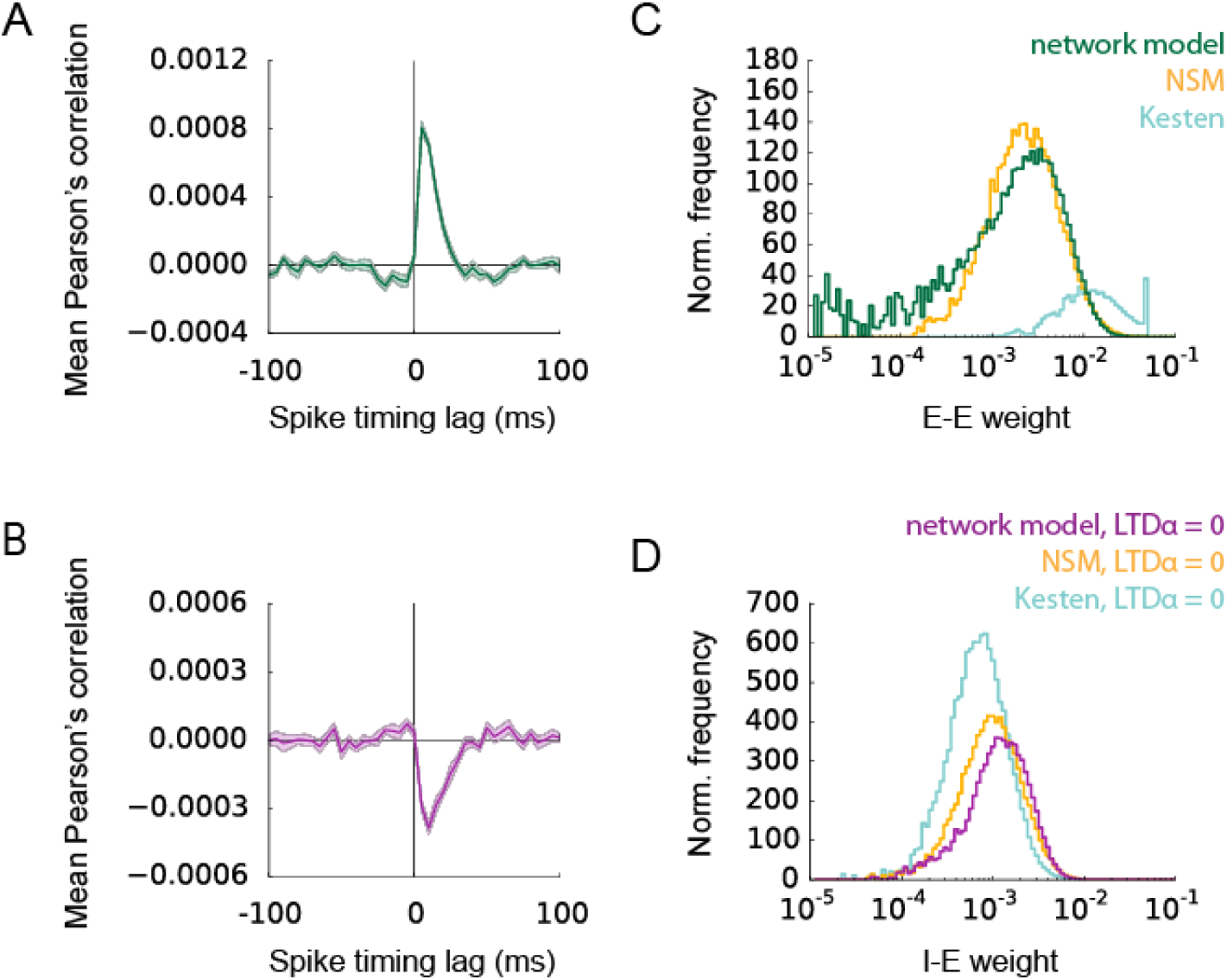
Comparison of E-E and I-E weight distributions to a nonuniform stochastic model (NSM) and a Kesten model. A: mean cross-correlations of spike times over all connected excitatory-to-excitatory neuron pairs. The mean correlation over 10 trials is shown in a solid line, while the shaded area indicates the standard error of the mean over 10 trials. Cross-correlations are recorded over a 200 second interval after the weights have reached equilibrium, with a spike bin of 5 ms. B: As in A but for all connected inhibitory-to-excitatory neuron pairs. C: weight distributions of E-E weights from the network simulation (green) and from two stochastic processes, the NSM (orange) and the Kesten model (blue). The distributions are averaged over 10 trials. D: as in C but for I-E weights. The distribution of the I-E weights is shown (purple) as well as the corresponding distribution from the NSM (orange) and the Kesten model (blue). Here, LTD_*α*_ = 0.

One may argue that the lognormal-like weight distributions obtained in the NSM are simply a result of the sampling of different numbers of STDP updates for different neurons from another lognormal distribution, but even if the firing rates are equal for all inhibitory and for all excitatory neurons in the network, the distributions of weights are still close to lognormal (Suppl. Fig. 3A-B, purple and green curves), as are the corresponding NSMs (Suppl. Fig. 3A-B, orange curves).

Since the NSM uses the spike cross-correlations to bias sampling from the iSTDP and eSTDP windows, we also wish to ask whether these correlations are strictly necessary to obtain the logormal-like distribution from STDP sampling. We therefore consider uniform sampling of the STDP window in the USM. The resulting distribution is similar to the distribution from the network simulation for I-E weights, but not for E-E weights (Suppl. Fig. 3C-D). Since the main difference between eSTDP and iSTDP in our model is the presence of LTD for eSTDP, we verified the distributions resulting from the USM and the NSM with various amounts of LTD (Suppl. Fig. 3E-F) . The distribution for the weights from the USM is only lognormal-like for zero LTD (Suppl. Fig. 3E) This explains why the USM matches well with the I-E weights from the network, since the iSTDP window contains no LTD for LTD_*α*_ = 0. In other words, since the USM is only close to lognormal when there is no LTD, this suggests the precise-spike time correlations do not matter in the absence of LTD. Implementing different iSTDP window shapes with LTD_*α*_ = 0 in the network model confirms this (Suppl. Fig. 4). Conversely, the USM does not resemble the distribution from the network when LTD is 80 percent of LTP or larger (Suppl. Fig. 3E). The NSM however yields a lognormal-like distribution regardless of the amount of LTD (Suppl. Fig. 3F). Since the peak in cross-correlation lies in the range for LTP in eSTDP, the spike correlations sample the LTP area of the STDP window more than the LTD area. Therefore, when the STDP window contains LTD, the presence of specific spike time correlations induced by the synapse is necessary to ensure a bias toward LTP and the emergence of a lognormal-like distribution of the weights. On the other hand, when there is only potentiation in the STDP window, random sampling of LTP events combined with SN already leads to a lognormal-like distribution.

### Lognormal-like distributions persist with spontaneous fluctuations

Recent results in cell cultures have shown that besides activity-dependent plasticity, spine sizes exhibit random changes that are independent of neuronal activity in the network [6] and of the local activity near each synapse [17]. In fact, [17] have demonstrated that the contribution of the spontaneous changes is at least as large as the contribution from the activity-dependent changes. Moreover, synapses continue to change in the presence of TTX, which abolishes spiking activity [6,23].

To see whether the weight distributions we found can still exist in the presence of such strong random fluctuations in the weight, we add independent changes to each weight (“weight noise”) throughout the simulation. The weight noise is sampled from a Gaussian distribution for both E-E and I-E weights. When the means of the Gaussians *μ*_we_ and *μ*_wi_ are positive, thereby favouring random potentiation in the weights, limiting distributions with lognormal-like shapes are found for both E-E and I-E weights (Fig. 10A-B). Importantly, these distributions persist in the absence of activity and activity-dependent changes (Fig. 10A-B, light coloured curves relating to TTX), in line with experimental findings [6,23]. However, when *μ*_we_ and *μ*_wi_ are negative, weights move away from their lognormal-like shape (Suppl. Fig. 5). Similarly to our results in Fig. 5C (blue curve), this suggests that if random fluctuations mostly consist of negative changes, the combination of random weight changes and multiplicative SN does not lead to lognormal-like weight distributions. This is equally valid in the absence of activity (Suppl. Fig. 5, TTX). We conclude that although dynamics and distributions of weights are affected, approximately long-tailed right-skewed distributions of weights can exist even in the presence of strong spontaneous fluctuations of E-E and I-E weights, as long as they contain a bias towards positive changes. Importantly, lognormal-like distributions can be seen even if the spontaneous fluctuations are of the same order of magnitude as the changes through LTP in eSTDP and iSTDP, and in the absence of activity-dependent plasticity like STDP.

**Fig 10.**
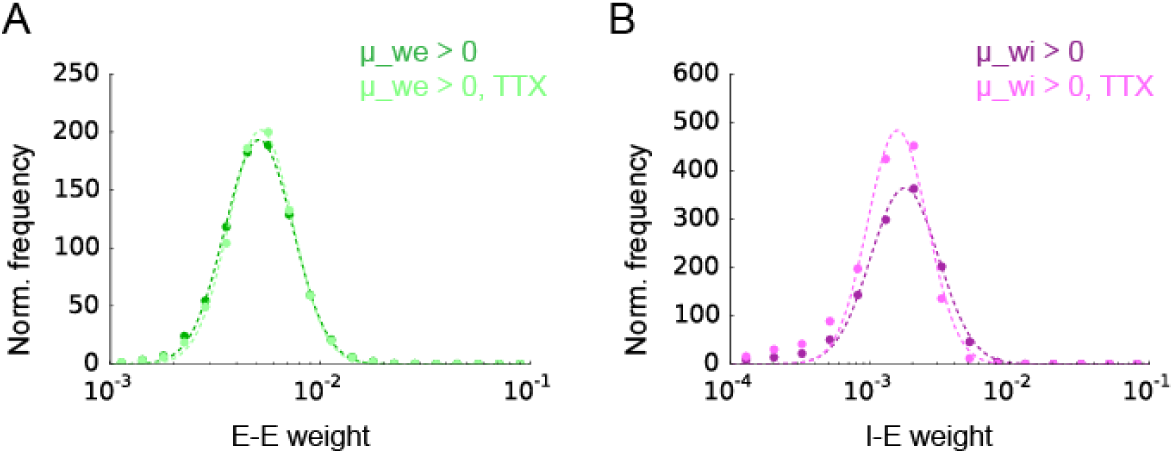
Weight distributions under spontaneous fluctuations of E-E and I-E weights under normal and silent conditions. A: Distributions of E-E weights in the presence of spontaneous fluctuations of E-E and I-E weights. In the case of TTX, weights change due to spontaneous fluctuations and SN only, meaning eSTDP and iSTDP are inactive. Dotted lines show the lognormal fit to the weight distributions. B: Same as in A but for I-E weights. In both panels, *μ*_we_ = *μ*_wi_ = 0.025 × 10^−3^ and *σ*_we_ = *σ*_wi_ = 0.1 × 10^−3^. Distributions are averaged over 10 trials.

## Discussion

The motivation of this work was to address whether cortical networks resemble a *democracy*, where the voice of every neuron has about the same weight, or whether they resemble an *oligarchy*, where a small set of neurons exerts an extreme influence over the network. Furthermore, we wanted to elucidate what mechanisms might give rise to such a concentration of power in cortical networks. Using a spiking neural network model we showed that a lognormal-like distributions of excitatory and inhibitory weights can result from the combination of additive STDP rules and multiplicative synaptic normalization as long as the net effect of STDP is biased towards potentiation. We also showed that these lognormal-like distributions can exist while individual weights fluctuate over time through either purely activity-dependent changes or spontaneous random alterations in the weights [17]. Moreover, we showed that small Gaussian variability in spiking thresholds results in highly skewed lognormal-like distributions of firing rates across the population [2] that are maintained over time [63]. The most active neurons in the tail of the distribution develop into *hub neurons* with strong efferent synapses, which potentiate due to a competitive advantage against synapses from less active neurons onto the same post-synaptic target neuron. Finally, we showed that the combination of high firing rate and strong efferent synapses allows these hub neurons to exert an extremely high influence on the rest of the network.

We found that lognormal-like distributions of excitatory and inhibitory synaptic efficacies appear in the network if the net effect of STDP is potentiating. Two factors contribute to this. First, the STDP windows favour potentiation. Specifically, our iSTDP window only contains contains potentiation. Second, the spiking correlations in the network bias the sampling of the STDP window such that potentiation dominates, as we have demonstrated for E-E synapses.

Another mechanism contributing to the lognormal-like distributions of synaptic efficacies is the presence of multiplicative normalization. While their is evidence for fast synaptic normalization mechanisms, keeping the sum of efficacies on dendritic branches approximately constant [43], we are not aware of evidence that such normalization is multiplicative. However, synaptic scaling mechanisms operating on longer time scales are thought to be multiplicative [9,45]. We consider it a plausible assumption that fast normalization also works multiplicatively [44]. A recent model by Tosi and Beggs [64] has also produced lognormal-like firing rates and synaptic efficacies. Their model relies on a hypothesized metaplasticity mechanism, for which experimental evidence seems to be lacking so far. In contrast, our model is capable of producing these distributions with only well-established plasticity mechanisms [15,26,28,43,45].

The key prediction from our model is the existence of a class of highly influential hub neurons that possess both high firing rates and strong efferent connections. This prediction could be tested using modern connectomics approaches in the following way. First, highly active neurons are identified through an appropriate recording technique. Second, the axon is reconstructed and the sizes of efferent synapses are measured. We predict that in such an experiment, highly active neurons will have stronger efferent synapses on average. The presence of strongly interconnected neurons in terms of synapse numbers has a strong effect on network functioning, affecting neuronal synchrony [65] and criticality [66]. Subgroups of strongly interconnected hub neurons have also been implicated in pathological states, namely hyperexcitability in the epileptic hippocampus [67]. It must be noted that in our study, the hub neurons do not possess on average larger numbers of synapses than the remainder of the neurons, since the network is connected randomly. However the hub neurons do have stronger outgoing weights and their impact on the network is highly pronounced, which may parallel the functional properties of neurons with more numerous synapses. The neural oligarchy that emerges from local processes at the neuron level may therefore be a critical feature of cortical functioning in health and disease.

The current network model is strongly simplified in a number of aspects. First, the synapses onto inhibitory neurons are not modified by plasticity, while experimental results in the visual cortex suggest these synapses do exhibit activity-dependent changes [68]. It will be necessary to address fully plastic networks to study the co-evolution of all synapses in an excitatory-inhibitory recurrent network. Moreover, although normalisation is implemented for each second in the current model, there is evidence for homeostatic mechanisms that come into effect over much longer periods [69]. We do predict that as long as the plasticity events in between the normalisation events are mostly instances of potentiation, and of small amplitude, the lognormal-like distribution of weights is guaranteed even over long timescales of normalisation. We also do not address here the pruning and creation of synaptic connections with multi-synapse connections per neuron pair [70–72]. How the distributions of inhibitory weights are affected by the appearance of multi-synapse connections is a subject for future study. Also, eSTDP and iSTDP are assumed to act equally fast, something that is problematic if inhibition itself acts as the major homeostatic mechanism by [69]. However if one assumes that synaptic normalisation in excitatory synapses acts rapidly, [43], the network need not rely on inhibitory plasticity to prevent runaway excitation. Although a comprehensive theoretical foundation for fully understanding the effects of inhibitory plasticity in recurrent neuronal networks may still be lacking [73], the various possible functions and consequences of iSTDP are beginning to be elucidated [74].

## Conclusion

In conclusion, our model suggests that cortical networks may resemble oligarchies, where a few neurons with high firing rates and strong efferent connections may exert a powerful influence over the rest of the network. On the one hand, this may have important implications for information processing in these networks [12,13]. On the other hand, this may also be important for network disorders such as epilepsy, where such highly influential neurons are believed to play a key role in the initiation and spreading of aberrant activity [67].

## Acknowledgments

JT is supported by the Quandt foundation. The authors would like to thank Felix Z. Hoffmann, Daniel Miner, Bruno Del Papa and Julijana Giorgjieva for helpful discussions.

## Supporting information

**S6 Video. The distribution of E-E weights evolves over time.** The distribution of E-E weights from an example trial is shown in the video, along with the lognormal fit to the distribution.The evolution is shown for 5-second steps over a 200 second interval. Weights are recorded just before SN.

**S7 Video. The distribution of I-E weights evolves over time.** The distribution of I-E weights from an example trial is shown in the video, along with the lognormal fit to the distribution.The evolution is shown for 1-second steps over a 40 second interval. Weights are recorded just before SN.

---

**Suppl. Fig. 1.**
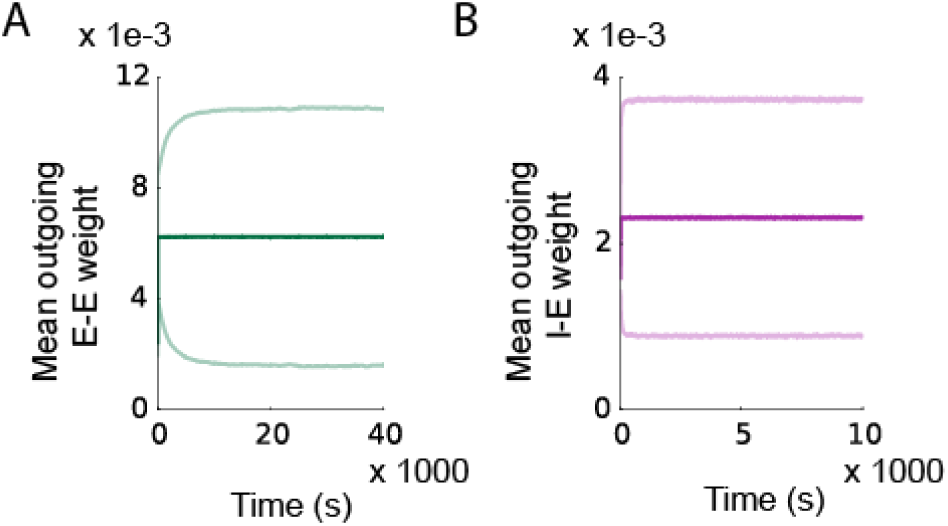
The weight dynamics stabilise over time. A: Evolution of the mean (dark green) and standard deviation (light green) of all E-E weights over time in a single trial. B: Evolution of the mean (dark magenta) and standard deviation (light magenta) of all I-E weights over time in a single trial.

**Suppl. Fig. 2.**
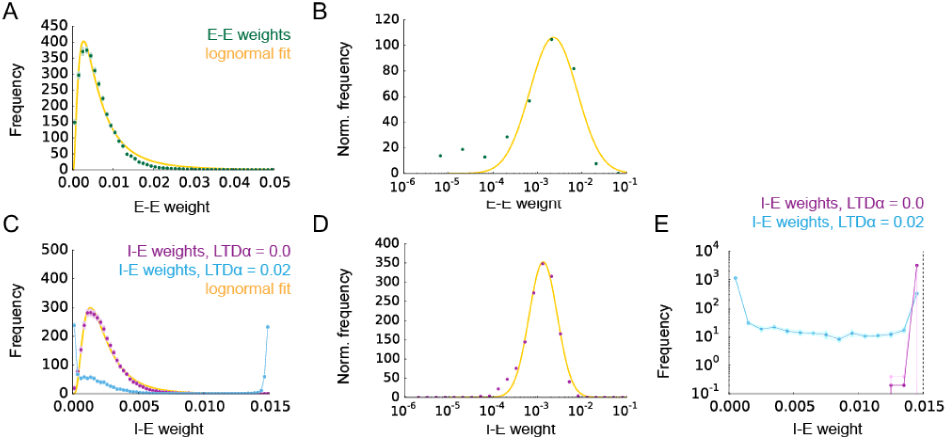
Average distributions of 10 trials of E-E and I-E weights, recorded just after SN. A: Mean distribution of E-E weights. The dark point show the average, the light point show the standard error of the mean over 10 independent trials. The orange curve is a lognormal fit to the mean distribution. B: Mean distribution of E-E weights plotted with uniform bin spacing on a logscale, in which bin values are corrected for their corresponding bin width. C: Mean distributions of I-E weights, for LTD_*α*_ = 0 (purple points), and LTD_*α*_ = 0.02 (blue points). A line is drawn between the blue points for visibility. D: As in B but for I-E weights, with LTD_*α*_ = 0. In all subfigures, the weights are recorded just after SN. E: Average distributions of I-E weights in the network without SN in the I-E synapses. The cases for LTD_*α*_ = 0 (purple), and LTD_*α*_ = 0.02 (blue) are shown. The vertical dotted line indicates the maximum I-E weight *T*_i,e_. The distributions are taken after 10000 seconds and averaged over 5 trials. The light colour shows the standard error of the mean. A line is drawn between the points for visibility.

**Suppl. Fig. 3.**
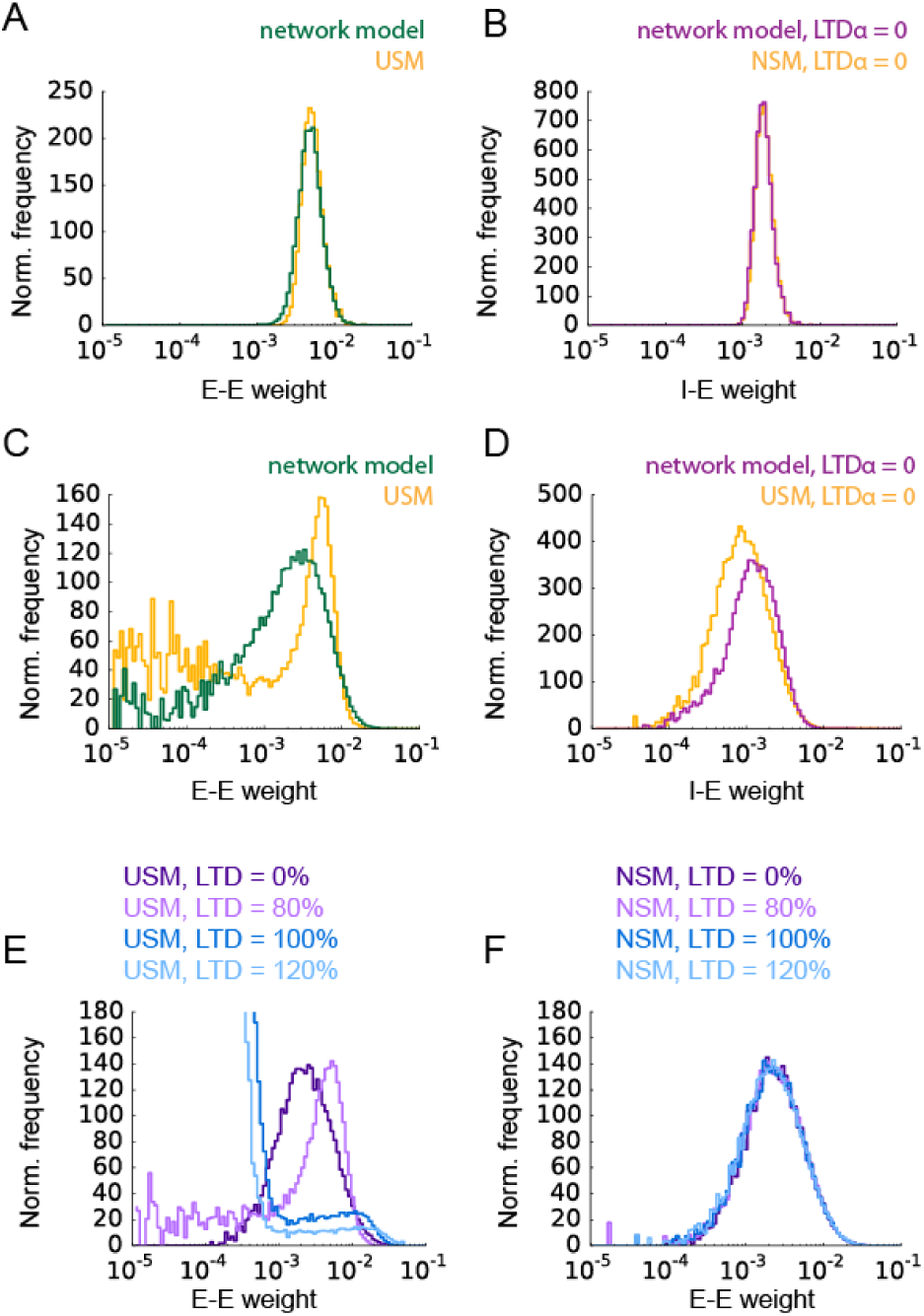
Comparison of E-E and I-E weight distributions from the network simulation to variations on the NSM and USM. A: The E-E weight distribution resulting from a network simulation with homogeneous firing rates for inhibitory and excitatory neurons (green) and its corresponding NSM distribution (orange) with eSTDP. B: as in A but for I-E weights and iSTDP, showing the distribution for I-E weights (purple) and its matching NSM distribution (orange). Here, LTD_*α*_ = 0. C: The E-E weight distribution from the network simulation (green) and its matching USM distribution (orange). D: The I-E weight distribution from the network simulation (purple) and its matching USM distribution (orange). E: The USM for E-E weights is shown for various quantities of LTD in the eSTDP window. F: as in E but for the NSM. All distributions are averaged over 10 trials.

**Suppl. Fig. 4.**
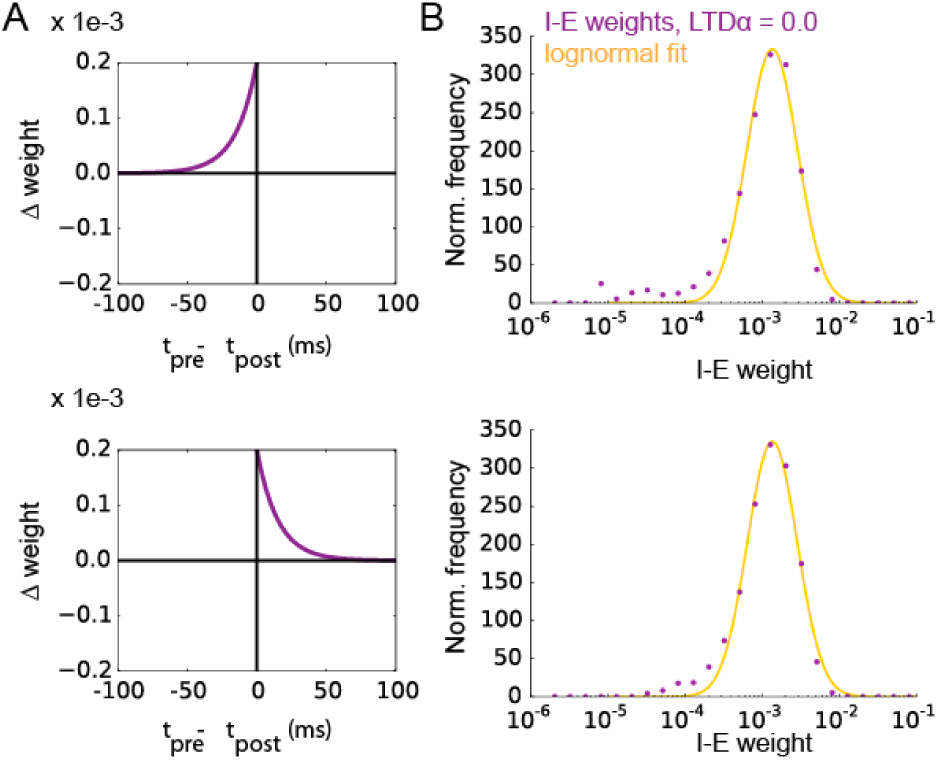
Average distributions of 10 trials of I-E and E-E weights in the network with iSTDP with SN in I-E synapses, shown for other iSTDP window types. A: iSTDP learning window and average I-E weights for pre-LTP iSTDP. The total amount of LTP is equal to the symmetric iSTDP window in Fig. 1. The yellow curve shows the lognormal fit. B: Same as in A but for post-LTP iSTDP.

**Suppl. Fig. 5.**
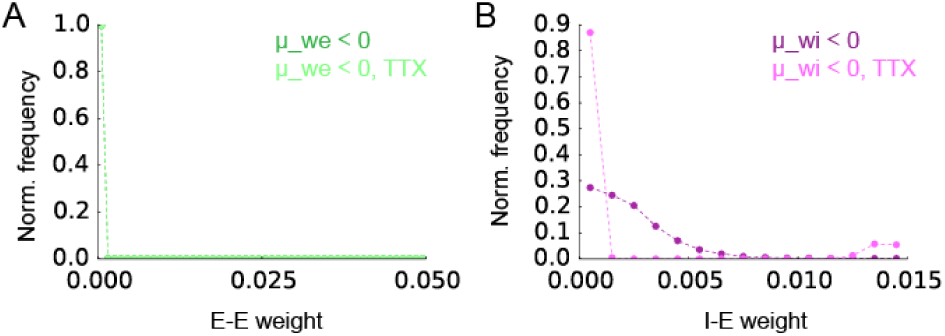
Weight distributions under spontaneous fluctuations of E-E and I-E weights under normal and silent conditions, for negative *μ*_we_ and *μ*_wi_. A: Distributions of E-E weights in the presence of spontaneous fluctuations of E-E and I-E weights. In the case of TTX, weights change due to spontaneous fluctuations and SN only, meaning eSTDP and iSTDP are inactive. The area of the histogram is normalised to 1.0. Dotted lines connect the data points for visibility. B: Same as in A but for I-E weights. In both panels, *μ*_we_ = *μ*_wi_ = –0.025 × 10^−3^ and *σ*_we_ = *σ*_wi_ = 0.1 × 10^−3^. Distributions are averaged over 10 trials, and each bin value is divided by the bin width.

